# Rapid and In-situ Detection of Biological Agents via Near-Infrared Spectroscopy and Cloud-Powered Neural Networks

**DOI:** 10.1101/2024.11.15.623740

**Authors:** Chloé Leseigneur, Isabelle Radgen-Morvant, César Metzger, Pierre Esseiva, Daniel Croll, Pascal Miéville

**Author notes:** Communication address.

## Abstract

The contagious nature of certain biological agents and the difficulty in treating infections render them a significant threat to public health and safety. In situations involving chemical, biological, radiological, and nuclear (CBRN) agents, effective detection is of paramount importance to prevent the undetected spread of these agents and enable swift and targeted responses tailored to the specific threat. Although portable detection tools are effective for chemical and nuclear detection, current biological detection methods face several challenges, including limited mobility, extended processing times, and varying accuracy. In the context of biological threats, where agents such as anthrax can be rapidly dispersed due to environmental factors and human activities, rapid detection is of paramount importance. It is imperative to develop field-applicable detection devices that are highly selective, capable of differentiating between biological and non-biological agents, as well as benign and harmful microorganisms. This study examines the potential of near-infrared (NIR) spectroscopy in conjunction with machine learning as a rapid in-situ biological detection method. The objective is to distinguish between biological agents and common white powders that are used as confounding agents in suspect letters. The non-pathogenic surrogates employed are safe and representative of typical biological warfare agents. The near-infrared (NIR) spectra of lyophilized bacterial and fungal surrogates, along with common white powders, were subjected to analysis and processing through the application of principal component analysis (PCA) and hierarchical clustering analysis (HCA). This resulted in the successful classification of the samples into distinct groups. The classification model demonstrated high accuracy in its prediction, thereby emphasizing the potential of the method for field detection of solid biological agents. These promising results suggest that NIR spectroscopy combined with machine learning could be further investigated as a rapid in-situ tool for biological detection in CBRN contexts.

## INTRODUCTION

CBRN (Chemical, Biological, Radiological and Nuclear) agents can be created intentionally or by accident, causing risks to human health, environment and animals.^1^ CBRN threats extends to accidents and attacks of varying scales and impacts, including those perpetrated by terrorists, pandemics and epizootics. Emergency responders, such as fire brigades, other specialized units (such as the EEVBS in Switzerland or the ATF in Germany) – and specialised military and civil protection/civil defence units, are responsible for CBRN protection and aim to prevent and reduce the impact of CBRN agents.^2^ CBRN detection technologies need to be practical, rugged and applicable in the field to identify threats and exclude as much as possible false positives. While all CBRN agents pose considerable risks, biological threats are associated with particular challenges, mostly because they can affect large groups and instil an atmosphere of fear and panic.^3^ Existing methods of detection are slow (>1h) and cannot be performed onsite as they require culture, and polymerase chain reaction (PCR). Biological threats require both emergency preparedness and technology development.

In the context of CBRN scenarios, *Bacillus anthracis*, the etiological agent of the anthrax disease, is recognized as a significant biological warfare agent in the 20^th^ century.^4^ Following the events of 9/11 in 2001, letters containing *Bacillus anthracis* spores were sent to various medias and Senate members, resulting in 11 cases of inhalation and five deaths.^5^ The anthrax powder was processed in order to to increase its propensity for aerosolization by adding electrostatic charges, reducing clumping, and uniformizing particle size to facilitate inhalation in a process named militarization.^6^ Since 2001, numerous anthrax fallacious threats involving harmless common white powder resembling flour, demonstrated the ease of fabricating such threats.^7^ Detecting anthrax spores in-situ remains challenging due to the need for rapid, accurate and reliable identification under high-stress conditions. The main difficulties in the detection of *Bacillus anthracis*, include its appearance as a harmless-looking powder, its potential for rapid dispersion and the necessity for prompt action to prevent widespread harm.^8^ Given its high relevance as a common biological threat to which CBRN units are potentially exposed,^9^ *Bacillus anthracis* was chosen as the target biological agent via non-pathogenic surrogates for this study on biological detection.

The most prevalent biological detection techniques encompass cell culture, polymerase chain reaction (PCR), enzyme-linked immunosorbent assay (ELISA), mass spectrometry, and biosensors. While cell culturing and plating are highly reliable, the necessity for several days to obtain results increases laboratory workloads and reliance on laboratory facilities.^10^ PCR technology, particularly real-time PCR, offers the potential for faster results within hours; however, it still requires laboratory settings and special equipment, which limits its field use.^11,12^ Similarly, ELISA also demands specific reagents and technical expertise.^13^ Mass spectrometry methods (e.g., MALDI-TOF) can accurately identify pathogen but are constrained by the need for complex equipment.^14^ Biosensors, on the other hand, offer a more rapid detection process; however, these technologies still require further development to achieve the necessary sensitivity and selectivity for reliable field use.^15,16^ Therefore, the principal limitation for the use of these techniques for CBRN response work is (a) their speed of detection/analysis (between 1 hour and many days depending on the method), (b) the lack of ruggedness of the equipment so that these cannot necessarily be transported and used in any field situation, and (c) their complexity of handling that may not be adapted for use by non-specialist first responder, military and civil defence/protection personnel.

Infrared spectroscopy is a method sensitive to chemical composition and architecture of molecules,^17^ applicable to biological systems, for example in bacterial strains identification.^18^ It is effectively used in the study of various biological materials, including lipids, proteins, peptides, bio-membranes, animal tissues, plants, clinical sample and microbial cells.^19^ The near-infrared (NIR) wavelength region of the infrared spectrum, extends from 13’000 to 4000 cm^-1^, covering a wave-length range from 769.23 nm ^1^ to 2500 nm.^19^ Discovered in 1800,^20^ NIR spectroscopy has been employed analytically since the 1950s.^21^ The method’s key advantages are a non-destructive and non-invasive technology.^22^ NIR technology has also been used for qualitative and quantitative control purposes in the pharmaceutical context ^23^ as well as for food sample analysis.^24^ Once calibrations is correctly performed, NIR becomes a relatively straightforward technique to use and can be made portable for terrain use.^25^ The device used in this study is the microNIR. A portable drug detection tool using NIR technology from VIAVI. The cloud-based software environment used to acquire the data is NIRLAB that promote the decentralization of forensic capabilities.^26^ NIRLAB identification is based on NIR spectra post-processing and neural network training. Similar methods have been used in previous studies using NIR spectroscopy for bacteria detection. For example, Kammies et al. report the analysis and differentiation of strains *B. cereus* and *E*.*coli, S*.*epidermidis* by hyperspectral NIR and PCR post treatment.^27^ Another study by Jiang et al. also demonstrated that PCA and HCA can effectively classify NIR spectra of different samples.^28^ Multiple studies have emphasized the effectiveness of combining NIR spectroscopy with machine learning algorithms, highlighting the utility of supervised methods such as neural networks and kNN in spectra analysis.^25,29,30^

The objective of this study is to assess the potential of portable near-infrared spectroscopy (NIR), as described above, in conjunction with cloud-based machine learning algorithms for the rapid and precise in-situ detection of biological agents of the bacillus type and discrimination with respect to commonly found white powders that may be used as fallacious threats and other biological agents such as fungi. In accordance with the preceding text, the *B. anthracis* has been selected as the target biological agent first due to its high importance in the CBRN context and second due to its compatibility with NIRLAB equipment, particularly its powdered weaponized form. In order to train the neural network and to ensure the safety of the experiment, non-pathogenic surrogates were selected based on their suitability for experimentation and their diversity. This selection allows for a comprehensive evaluation while avoiding the inherent dangers associated with highly pathogenic strains. The study encompasses an examination of common white powders, including baking soda, sugar, and talcum, which are pertinent in the context of anthrax fallacious threats.^31^ This approach aims to enhance the application of NIR spectroscopy in public safety and CBRN scenarios, ensuring rapid and reliable detection of potential biological threats in situ.

## METHODS

### Biological non-pathogenic surrogates

Twenty-three biological non-pathogenic surrogates were selected, comprising 11 bacteria (see Table 1) and 12 fungi (see Table 2). The selection of each surrogate was made with due consideration to their safety, while maintaining proximity to biological agents. In this context, proximity is understood in terms of biochemical structure for the bacteria and physical appearance for fungal powders. The objective was to obtain a diverse range of samples and to prepare them in the closest possible powdered form.

**Table 1.** List of bacteria used in the present study as b. anthracis non-pathogenic surrogates with comments on selection criteria. ^*^ classified by the U.S. CDC as category A bioterrorism agent, ^**^ classified by the U.S. CDC as category B bioterrorism agent ^42^.

| Bacteria | Comments | References |
| --- | --- | --- |
| *Bacillus cereus* | common soil and plant dwelling bacterium, spore-forming, can be aerosolized, opportunistic human pathogen (inter alia food safety threat), closely related to *Bacillus anthracis* * and *B. thuringiensis* | 32 |
| *Bacillus thuringiensis* | soil dwelling bacterium, common biocontrol agent, suspected opportunistic pathogen (rare occurrences), closely related to *Bacillus anthracis* * and *B. cereus* | 33,34 |
| *Bacillus amyloliquefaciens* | used as biocontrol agent in agriculture, synthesizes a narrow-spectrum natural antibiotic against *B. anthracis* | 35,36 |
| *Bacillus subtilis* | soil and gastrointestinal (e.g., humans, ruminants) bacterium, model species for the *Bacillus* genus, important industrial organism | 37 |
| *Escherichia coli* | normal part of the gastrointestinal microflora of humans, model bacterium, many pathotypes can cause disease in humans (food safety threat)** | 38 |
| *Pseudomonas putida* | soil dwelling bacterium, used as biocontrol agent in agriculture, used in bioremediation, opportunistic human pathogen | 39 |
| *Enterococcus faecalis* | normal part of the gastrointestinal microflora of humans, opportunistic human pathogen | 40 |
| *Staphylococcus epidermidis* | normal part of the skin microbiota of humans, opportunistic pathogen, closely related to human pathogen *Staphylococcus aureus* (of note: the CDC classifies staphylococcal enterotoxin B as a category B bioterrorism agent, other staphylococcal toxins are common causative agents of food poisoning) | 41,42 |
| *Paracoccus* spp. | i.e. *Paracoccus yeei*, opportunistic human pathogen (rare accounts) | 43 |
| *Burkholderia terrae* | soil dwelling bacterium, non-pathogenic, sister species to pathogenic *Burkholderia mallei* ** and *Burkholderia pseudomallei* ** |  |
| *Lactobacillus rhamnosus* | the *Lactobacillus* genus has been studied with NIR as food contaminant | 44 |

**Table 2.** List of fungi used in the current study as non-bacterial biological b. anthracis surrogates with comments on selection criteria.

| Fungi | Comments | References |
| --- | --- | --- |
| *Saccharomyces cerevisiae* | model yeast species, opportunistic human pathogen | 45 |
| *Gliocladium catenulatum* | used as biocontrol agent in agriculture | 46 |
| *Ophiocordyceps heteropoda* | known instances of human poisoning cases | 47 |
| *Elaphocordyceps subsessilis* | soil fungus known to naturally produce ciclosporin A | 48,49 |
| *Candida albicans* | prevalent opportunistic human pathogen | 50 |
| *Trichoderma spp.* | soil dwelling fungi, several strains are used as biocontrol agents in agriculture, *Trichoderma longibrachiatum* (common house mold) can be toxic to humans, produces mycotoxins | 51 |
| *Clonostachys rosea* | used as biocontrol agent in agriculture | 52 |
| *Fusarium graminearum* | soil dwelling fungi, produces mycotoxins | 51 |
| *Aspergillus niger* | opportunistic human pathogen, related to the human pathogen *Aspergillus fumigatus* | 53 |
| *Exophiala xenobiotica* | an opportunistic black yeast, *Exophiala* is one of the rare fungi causing infection in immunocompromised people | 54 |
| *Penicillium commune* | common fungi contaminating cheese, it also produces mycotoxins | 55,56 |
| *Alternaria spp.* | can be plant pathogens, producing mycotoxins implicated in mammals' cancer, it has potential of emergence as a human pathogen | 57 |

All strains mentioned in Tables 1 and 2 were obtained from the Microbiology laboratory collection at the University of Neuchâtel. All bacteria were cultivated on agar plates, from which colonies were extracted and inoculated into a nutrient broth solution of 200ml. The same process was followed for fungi except they were inoculated into malt broth. The culturing process required different times for each biological species. The suspensions were distributed into four vials of 50ml, which were then centrifuged at 900 rpm for three minutes at 8600 g. Subsequently, the samples were lyophilized at -50°C and 0.05 mbar for 24 hours. Following lyophilisation, all specimens were crushed with a mortar prior to analysis. To compare to the biological spectra, 21 common white powders were chosen. The common white powders represent typical anthrax fallacious threats, mostly easy to access. The common white powders can be classified into four categories (Table 3.).

**Table 3.** Samples defined in the text as common white powder, mostly easily accessible, corresponding in the current study as non-biological fallacious threats encountered by CBRN responders.

| Categories | Powders | Formula | Comments |
| --- | --- | --- | --- |
| Cooking | Crystalized sugar | $C_{12}H_{22}O_{11}$ | |
| Cooking | Cornstarch | $(C_6H_{10}O_5)_n$ | Sugar polymer |
| Cooking | Kitchen salt | NaCl |  |
| Salts | Sodium carbonate | $Na_2CO_3$ | |
| Salts | Sodium bicarbonate | $NaHCO_3$ | |
| Salts | Potassium carbonate | $K_2CO_3$ | |
| Salts | Dipotassium phosphate | $K_2HPO_4$ | |
| Salts | Disodium hydrogen phosphate | $Na_2HPO_4$ | |
| Salts | Potassium dihydrogen phosphate anhydrous | $KH_2PO_4$ | Monopotassium phosphate |
| Salts | Magnesium chloride anhydrous | $MgCl_2$ | |
| Salts | Ammonium chloride | $NH_4Cl$ | |
| Household and industrial products | Talcum | $Mg_3Si_4O_{10}(OH)_2$ | |
| Household and industrial products | Calcium carbonate | $CaCO_3$ | Sampled from chalk |
| Household and industrial products | Anhydric citric acid | $C_6H_8O_7$ | |
| Household and industrial products | Celite |  | Diatomaceous earth |
| Medicines and supplements | Aspirin |  | Acetylsalicylic acid |
| Medicines and supplements | Neocitran |  | Medicine for cold and flu symptoms |
| Medicines and supplements | L-isoleucine |  | Amino acid |
| Medicines and supplements | L-menthol |  |  |
| Medicines and supplements | L-proline |  | Amino acid |
| Medicines and supplements | Propylparaben |  | Preservative |
| Medicines and supplements | Ethylparaben |  | Preservative |

### NIR Analysis

The MicroNIR Onsite W 1700 from VIAVI was employed as a portable near-infrared (NIR) tool, connected to the NIRLAB Mobile application via Bluetooth.^26^ The system operates within the spectral region spanning 950 to 1650 nm. The biological samples (Tables 1 and 2) were lyophilised using the Lyoquest Telstar laboratory freeze dryer, while non-biological common white powders (Table 3) were crushed with a mortar in the case of grains (e.g., chalk) and directly sampled. The resulting lyophilised powders were weighed and transferred to aluminum plates (Figure 1) with a spatula. Aluminum plates are optimal to minimize reaction with the sample, interference, or beam absorption during the analysis with the microNIR device. The samples were arranged in a manner that ensured coverage of the device’s beam, thereby allowing for an effective analysis.

**Figure 1.**
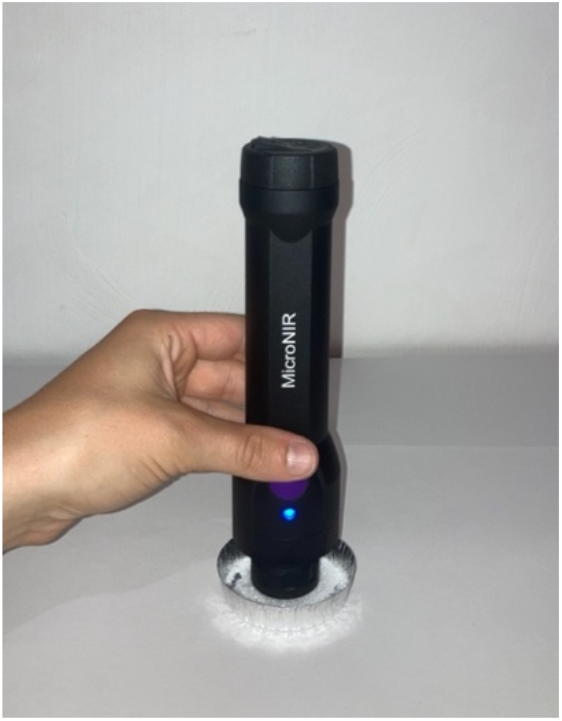
The microNIR device used to analyse all samples on a typical aluminium plate. In this specific example, the sample is crystalized sugar.

The microNIR device was positioned in direct contact with the samples. Upon activation of the microNIR device, the resulting spectra were promptly recorded in the NIRLAB application. Each substance was subjected to three separate samplings and each sample was subjected to three replicate analyses for signal averaging, resulting in a total of nine spectra per powder. On some samples, depending on the acquired spectra quality, a fourth replicate analysis was performed.

### Data processing and neural-network training

The combination of the sampling and measurement replication method described above with the list of samples (Tables 1, 2 and 3) yielded a total of 471 spectra, which were exported in Excel format (.csv) to Orange Data Mining software for subsequent data analysis. In order to identify a fingerprint of the substance with NIR, it is necessary to reduce the noise and increase the signal from the chemical information. This can be achieved through the application of data pre-processing. The pre-processing methods employed were the standard normal variate (SNV) to normalise the spectra and the Savitzky-Golay filter with and without a second derivative and a second-order polynomial. The selected spectral range was trimmed and retained between 950 and 1641.8 nm. The initial analysis was conducted using principal component analysis (PCA), with the objective of identifying any inconsistencies or outliers present in the data set.^25^ PCA method enables the reduction of the dimensionality of multivariate data,^52^ while ensuring the maintenance of its intrinsic information content. This method accounts for the majority of the data variance. A preliminary PCA was conducted on all spectra. In order to account for the intra-variability within a given group (species or substance), an average spectrum is calculated for each fungal and bacterial species, as well as for each non-biological common white powder. Hierarchical clustering analysis (HCA) is a valuable method for organizing samples based on their pairwise similarity.^58^ The similarity between the samples was assessed using the HCA method, with the cosine distance measure employed as the metric. The pre-processed samples of all spectra were subjected to HCA. A distance map (heat map), based on the averaged spectra, was constructed to facilitate the visualization of clusters within the data set. The samples were randomized and divided into two groups: a calibration group comprising two-thirds of the samples and a prediction group comprising the remaining third. To assess the efficacy of the classification models, a series of machine learning algorithms, including neural networks, support vector machines (SVM), and k-nearest neighbors (kNN), were employed. The performance of these models was evaluated using a range of performance metrics, including the area under the receiver operating characteristic curve (AUC), classification accuracy (CA), F1 score, precision, recall, and the Matthews correlation coefficient (MCC). This comprehensive approach ensured a thorough evaluation of the models’ performance.^59^ A confusion matrix was constructed for the purpose of identifying specific instances that had been misclassified and of ascertaining the proportion of instances that had been predicted. The confusion matrix elucidates the patterns of errors inherent to the models.

## RESULTS

As previously stated, a total of 471 spectra were acquired. Figure 2 depicts the NIR spectra after standard normal variate (SNV) and second derivative pre-processing, which demonstrate considerable chemical variability, particularly evident in the talcum samples, which exhibit elevated peaks at approximately 1370 and 1390 nm. The spectra of bacteria and fungi display greater clustering, with analogous peaks observed within each group.

**Figure 2.**
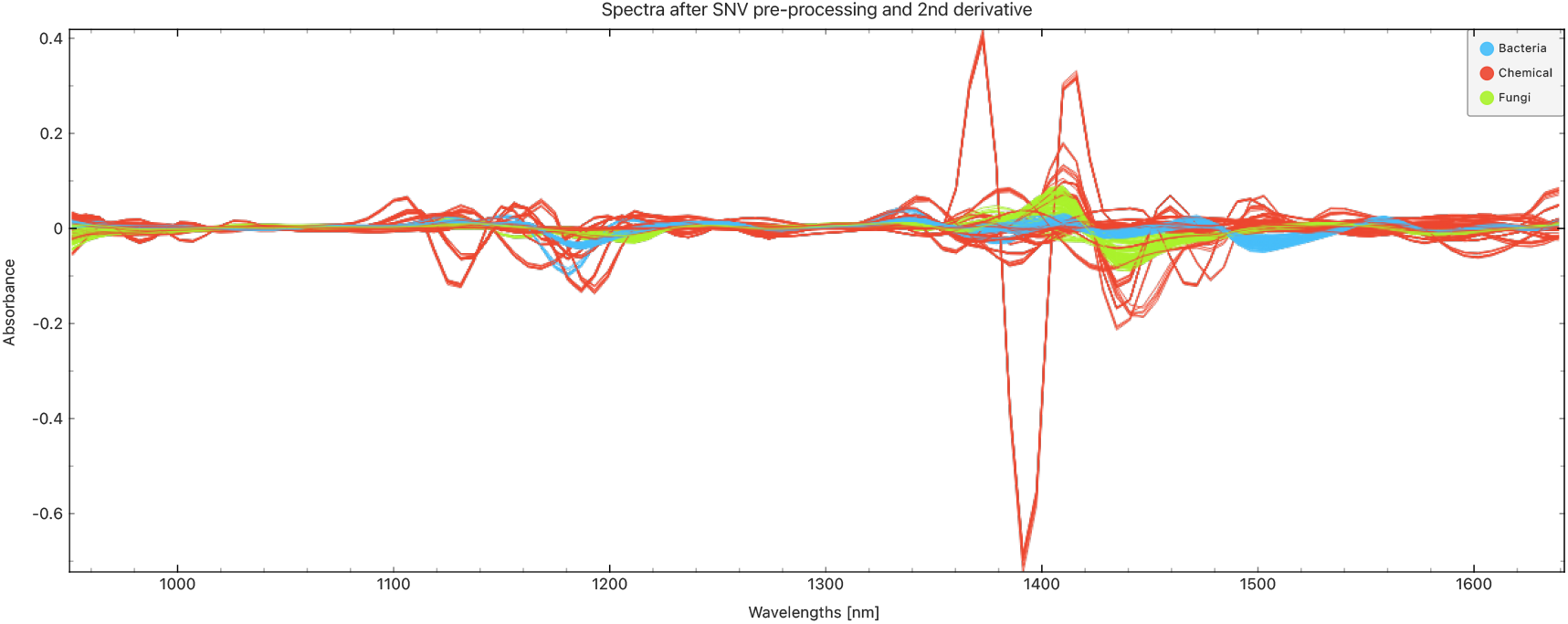
NIR spectra of bacteria (blue), fungi (green) and common white powders (Chemical, red) after SNV and second derivative pre-processing.

A PCA was conducted on all aforementioned spectra following pre-processing to reduce the dimensionality of the dataset while retaining the most significant variance. This enabled the discovery of patterns and relationships that might otherwise have remained obscured in the high-dimensional spectral data. The resulting data are presented in Figure 3 for the reader’s convenience. A more segregative representation is provided by the PCA of the averaged spectra per sample. The averaging reduces the variability between individual samples and demonstrates a more pronounced separation between the groups (Figure 4). A clear segregation is evident in the PCA plot of mean spectra, with bacteria and fungi forming discrete clusters. However, the segregation is less pronounced regarding the common white powder substances, as several can be found within the fungal group, including cornstarch, sodium carbonate, sodium bicarbonate, and dipotassium phosphate. Furthermore, propylparaben was the only chemical substance found in the bacterial group.

**Figure 3.**
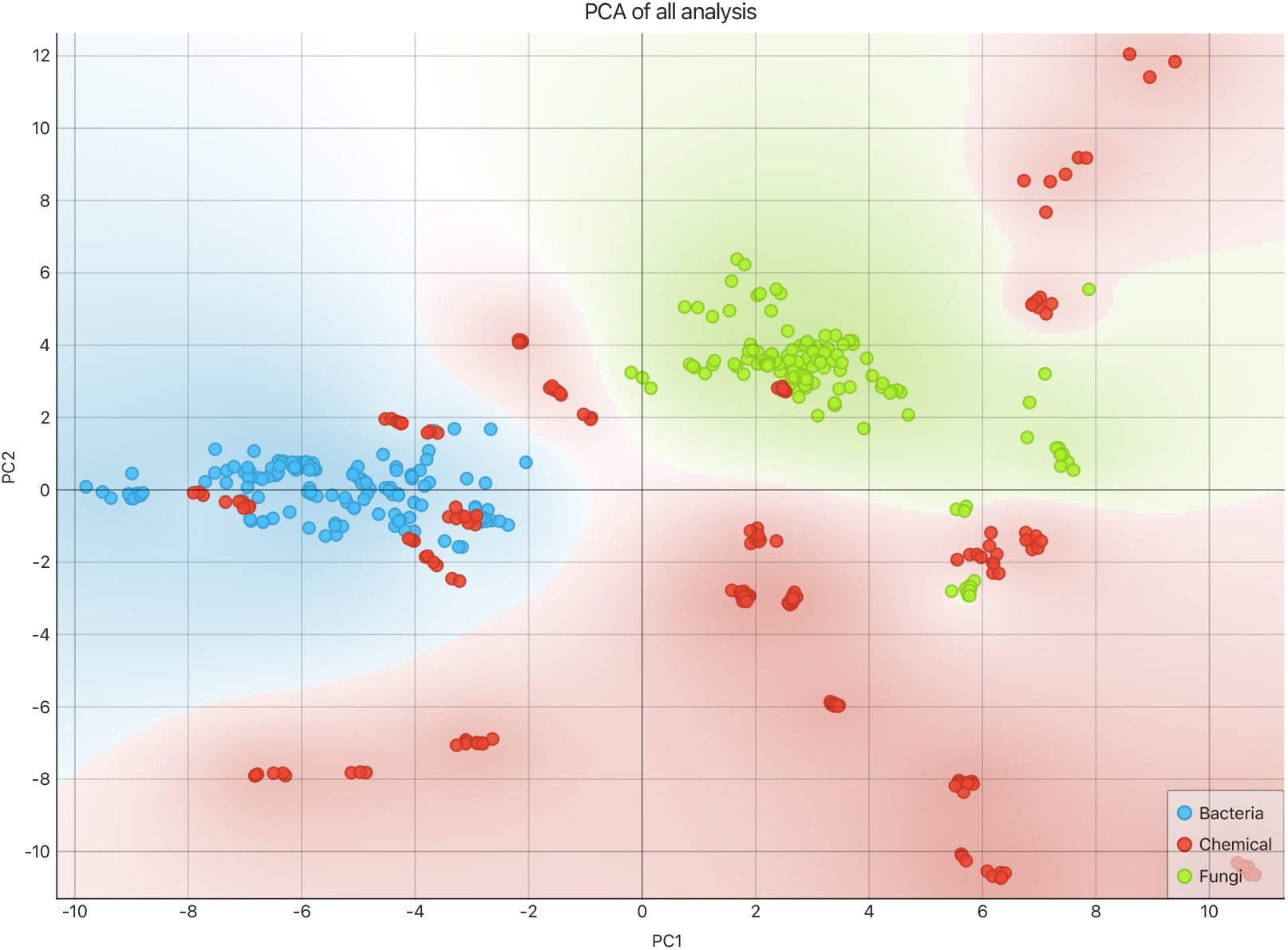
PCA of the 421 spectra collected on the samples listed in Tables 1 to 3, following the SNV and second derivative pre-processing as shown in Figure 2. The red zone represents the group of non-biological common white powders, the blue zone corresponds to the bacterial group of spectra, and the green zone corresponds to the fungal group.

**Figure 4.**
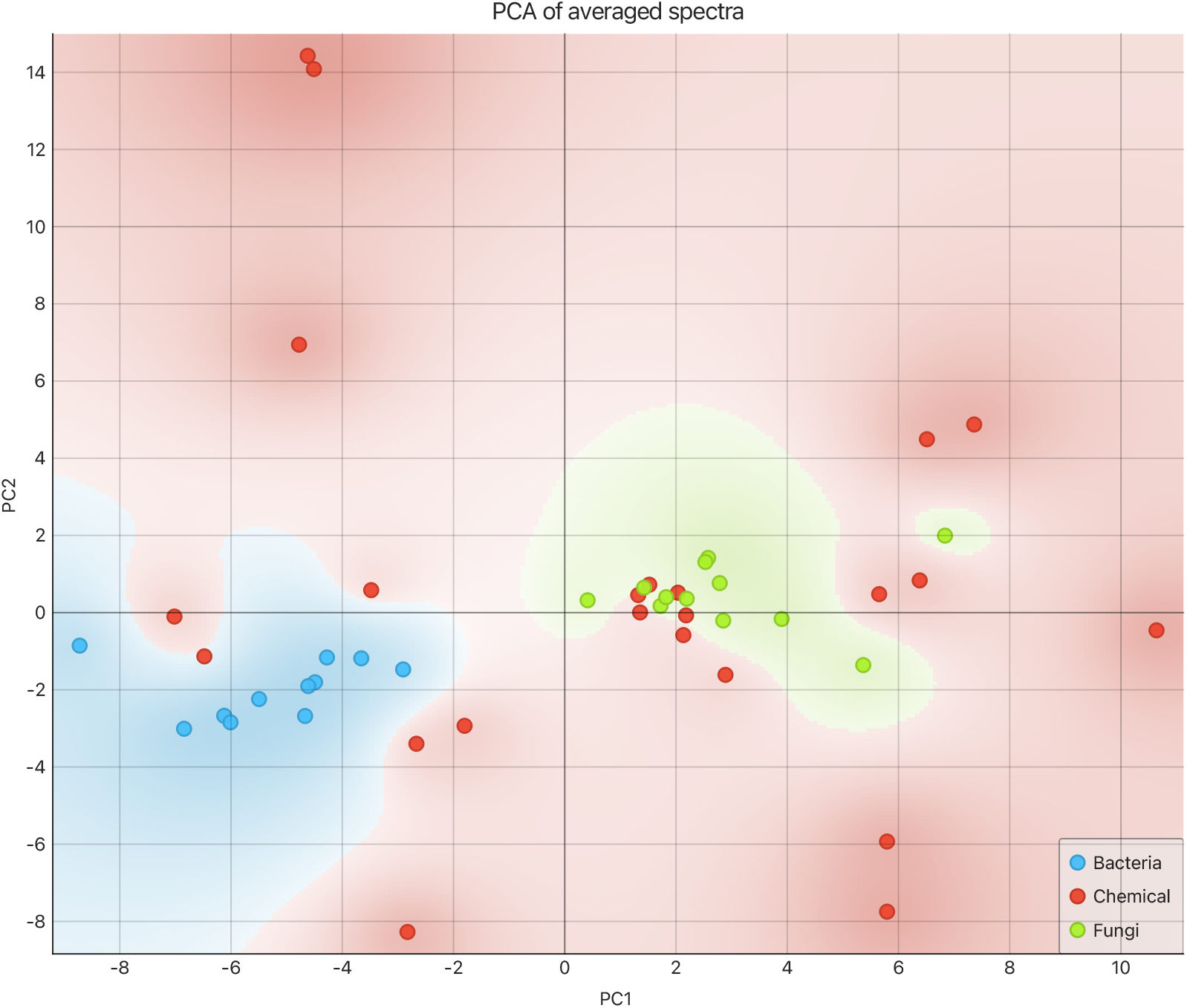
PCA of mean spectra of bacteria (blue), chemical (red) and fungi (green).

In order to evaluate the degree of spectral discrimination achieved through PCA, a hierarchical cluster analysis (HCA) was performed on the same set of 421 spectra. The aim is to conduct a comprehensive evaluation to ascertain whether the replicates within the dataset demonstrate discernible clustering patterns or whether there is a notable degree of overlap between the groups. The results of the HCA, presented as a distance map in Figure 5, indicate that bacterial species exhibit greater clustering tightness (see the main centered diagonal blue square), highlighting stronger within-group spectral similarity compared to other groups. It is notable that non-biological common white powder samples form discrete, smaller clusters than biological ones and are predominantly located in the lower half of the diagonal. Moreover, some common white powders, including magnesium chloride, ammonium chloride, and Neocitran, diverge from the primary common white powder clusters, forming smaller clusters at the top of the diagonal. This suggests that these compounds possess spectral characteristics analogous to those observed in other sample types. These observations are consistent with the PCA results, which identified two major bacterial and fungal clusters surrounded by greater variation in the common white powder spectra. Therefore, the combination of PCA and HCA provides a robust framework for evaluating the degree of spectral discrimination and clustering tendencies across the dataset.

**Figure 5.**
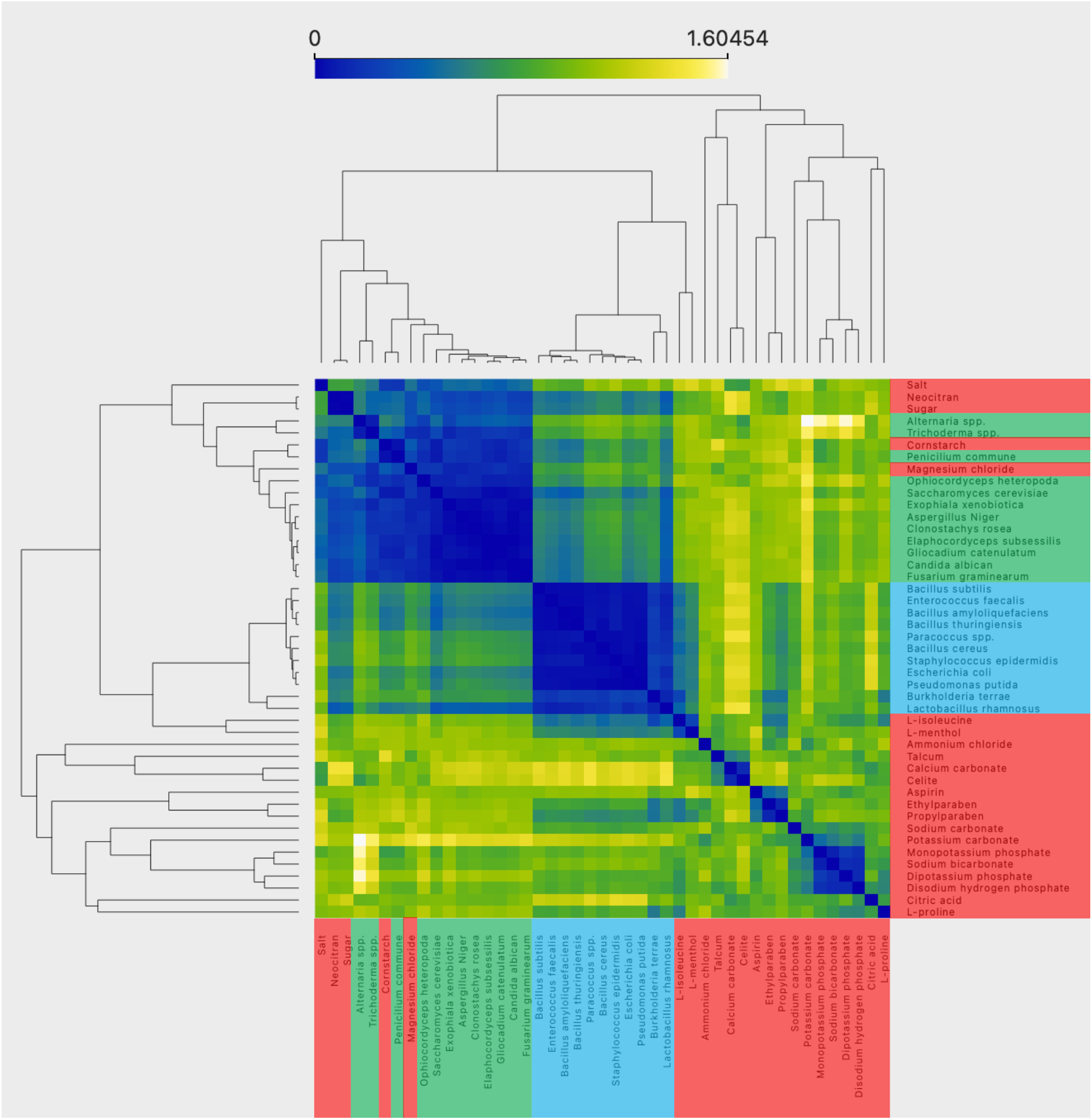
HCA with a distance map visualization. The mean spectra of the samples are used with cosine distance values, with dark blue representing a high degree of similarity and red, blue, and green representing progressively lower degrees of similarity. The legends have the same colors as the PCA, with red indicating chemicals, blue indicating bacteria, and green indicating fungi. The left top square is predominantly composed of fungal samples, while the middle square is composed of bacterial samples.

A model performance evaluation was conducted to test the robustness of the method. The performance of the classification of the dataset was evaluated using a neural network (NN), a support vector machine (SVM) and a k-nearest neighbour (kNN) model. The results demonstrated that all models exhibited strong performance. While SVM and kNN also performed well, the NN model achieved the best classification score across all metrics (Figure 6.a). The confusion matrix provided the proportion of instances between the predicted group and the calibration group for the NN model (Figure 6.b), confirming the accuracy of the chosen model.

**Figure 6.**
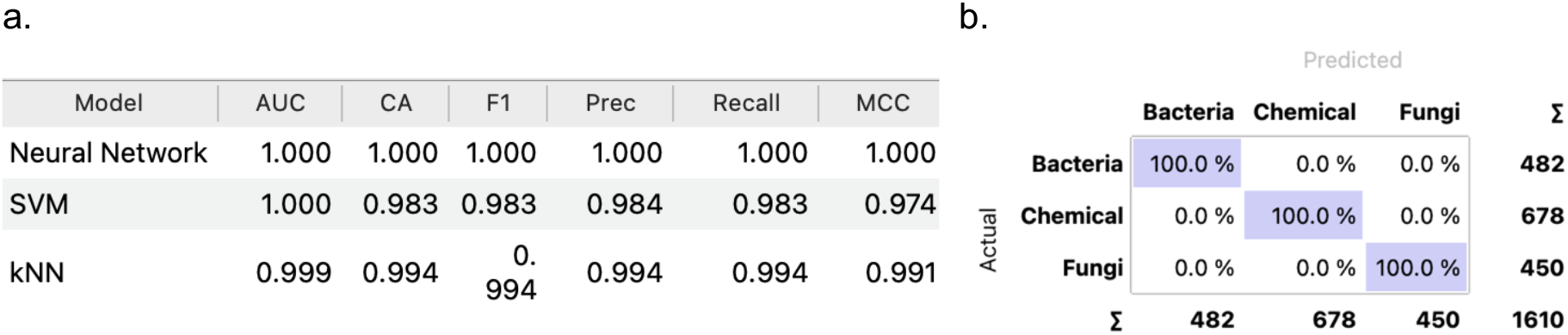
**a** This table presents the performance scores of each model, NN, SVM and kNN. The performance metrics used are area under ROC curve (AUC), classification accuracy (CA), F1 score, precision, recall and Matthews correlation coefficient (MCC). **6.b** The second table shows the confusion matrix of the NN model.

## DISCUSSION

The objective was to assess the potential of the portable near-infrared (NIR) spectrometer MicroNIR Onsite W 1700 from VIAVI, in conjunction with NIRLAB cloud-powered NIR spectroscopy (NN), as an in-situ rapid method for the detection of chemical, biological, radiological, and nuclear (CBRN) biological agents. This was achieved through the analysis of over 11 different lyophilised bacteria, 12 different lyophilised fungi, and 21 common, easily accessible white powders. In this context, lyophilised bacteria and fungi serve as non-pathogenic surrogates for typical militarised *B. anthracis*, allowing for the demonstration of the feasibility of rapid and significant discrimination between biological and non-biological common white powder groups of samples. The trained NN also demonstrates the possibility of automatically classifying a potential new powdery sample measured with the MicroNIR spectrometer with a high degree of confidence, confirming the possibility of using this apparatus in combination with the NIRLAB software to perform CBRN in-situ biological agent identification. It is necessary to provide a commentary on the 100% discrimination achieved on the NN (Figure 6.b). The quality of the spectra acquisition and subsequent classification is undoubtedly influenced by the specific characteristics of the samples, including their granularity and homogeneity. These attributes can be subject to variability due to the inherent unpredictability of the real-world conditions employed in the preparation of microorganisms or common white powders.^60^ For example, the lyophilisation, which usually preserves viability but can affect cellular structures and impact detection of surface protein,^61^ could impact the spectrum. It would also be relevant to explore the influence of the device temperature on the bacterial spectrum, as well as the operational temperature of the NIR devices in field conditions.^62^ Furthermore, historical instances of biological attacks utilising *B. anthracis* have involved the processing and treatment of the bacteria in a manner that was specifically designed to suit its intended use as a weapon.^6^ Such treatments could potentially impact the NIR analysis of the biological agent. Further experimentation with authentic *B. anthracis* and potentially other biological warfare agents is necessary to corroborate the findings presented in this study and to fully validate its use in operation.

The PCA facilitated the visualization of the differentiation between the biological groups, including bacteria and fungi. In addition, the analysis revealed a notable diversity of the common white powders. The results of the PCA visualization are promising and are corroborated by the HCA, which also reveals the existence of the three groups of common white powders. A number of the common white powders exhibit a closer proximity to the fungi than to the other common white powders. Among these substances, cornstarch, sugar, salt, and magnesium chloride are included in the fungi cluster. The fact that cornstarch is a polysaccharide, which is quite similar in structure to chitin, itself a polysaccharide and a primary component of the cell wall of fungi, may explain why this powder is related to fungi. The same explanation is likely also valid for the proximity of other sugars. As previously noted, the neural network model yielded optimal predictions for the three groups, substantiating its resilience and underscoring prospective avenues for advancement. Nevertheless, it is important to exercise caution, as a 100% classification results may indicate overfitting, especially considering the small size of the dataset. A larger dataset would confirm the high specificity and sensitivity of the method by reducing the risk of overfitting.^63^ These results nevertheless underscore the interest of advanced machine learning algorithms combined with portable NIR tools for rapid and accurate analysis of biological samples.

A further potential limitation of a spectral approach is that it does not separate substances and provides an average spectrum of the measured mixture. Consequently, the effect of an eventual matrix used for the culture or the dispersion of the biological agent can be significant.^26^ Further research could investigate the discrimination of a biological agent within a non-biological dispersion matrix and the combination of different biological strains. The developed method is dependent on the ability to recognize substances within an existing database. It is therefore unable to identify unknown types of molecules or substances for which the NN has not been trained. This highlights the necessity of creating a larger database of biological agents and potential fallacious threats. Furthermore, the focus of this study is on bacterial detection, which overlooks significant threats like viruses.

As previously stated, the objective of this study is to illustrate the potential of rapid and in situ differentiation between biological and non-biological samples to alert CBRN responders. The initial objective was not to identify a specific biological agent. Nevertheless, the demonstrated differentiation between fungi and bacteria and the ability to distinguish each bacterial strain on the PCA provide a promising foundation for exploring the potential of selective identification. This could prove a valuable addition to CBRN operations, and it may also have applications in the detection of pathogenic strains in other fields, including medical microbiology, food science, and forensic sciences. To extend the potential of the method, it would be advantageous to create a database of NIR spectra that involves the various stakeholders addressing all aforementioned domains. In this context, the cloud-based neural network would facilitate the rapid updating of information on pathogens, ensuring that the database remains current and accessible to a large group of actors and that the NN is continuously updated.

## CONCLUSION

This study demonstrates the potential of near-infrared portable spectroscopy in conjunction with cloud-based neural networks for expeditious, in-situ biological threat detection in contexts pertaining to chemical, biological, radiological, and nuclear (CBRN) incidents. The application of a cloud-based machine learning algorithm enables the effective classification of the studied groups and the discrimination of samples between biological and non-biological. It is proposed that NIR could serve as a rapid and simple-to-use biological detection device for biological CBRN emergencies. Moreover, the application of this technology could be extended to other fields, including microbiology, food analysis, and forensic science. In this context, the creation of a NIR spectra database of biological agents and non-biological potential fallacious threats involving the various stakeholders that address microbiological challenges would be beneficial.

## Acknoledgments

The authors acknowledge the ETH Domain for its support through the Forschungsinfrastrukturen Program. They would also like to acknowledge the complementary support offered by Cap. Bernard Tschopp and Col. Nicolas Schumacher of the “Service Incendie et Secours” of Geneva.The authors would like to thank Florentin Coppey of the University of Lausanne and NIRLAB founder for providing the microNIR device and support. We thank Prof. Pilar Junier and lab assistant Ilona Palmieri of the microbiology lab of the University of Neuchâtel for their assistance.

## Authors contributions

Conceptualization : P.M, C.L, C.M, P.E, I.R-M

Funding : P.M, D.C

Methodology : P.M, D.C, C.M, I.R-M

Resources : P.E, D.C

Software : P.E, I.R-M, C.L

Supervision: P.M, D.C Writing: C.L, P.M, D.C Revision: P.M, D.C

